# Dance experience shapes local brain dynamics and shared responses during naturalistic action observation

**DOI:** 10.64898/2026.11.15.766898

**Authors:** Da Chang, Donghui Song, Henyi Rao

**Affiliations:** College of Education Science, Hengyang Normal University, Hengyang, China; Department of Psychology, University of Science and Technology of China, Hefei, China

**Keywords:** action observation, dance expertise, brain entropy, intersubject correlation, coactivation patterns, naturalistic fMRI

## Abstract

Skilled action observation unfolds over time: viewers follow changing movements, coordinate what they see with prior experience, and form a response to the performance as a whole. Dance expertise is known to alter action-related activation and the similarity of spectators’ responses, but its association with local BOLD dynamics during an extended performance remains unclear. We reanalyzed fMRI from 14 professional dancers and 11 novices watching the same 33-minute dance film; one choreographer provided a separate reference. In the first 678 volumes, we measured voxelwise brain entropy (BEN) using sample entropy before summarizing it in Schaefer 400 parcels and seven networks, calculated windowed entropy from parcel-mean signals, and examined intersubject correlation (ISC) and four group-blind coactivation patterns (CAPs). After adjustment for local gray-matter volume, age, sex, and motion, dancers had higher static BEN in 32 parcels and six networks (FDR q < .05). Dynamic BEN was higher in control and default-mode networks. Dancer-dancer ISC exceeded novice-novice ISC in five networks. Dancers occupied one common CAP less often and another more often, while performer synchrony and acceleration tracked state occurrence across spectators. These converging observations show that dance experience is expressed in local temporal irregularity, shared spectator responses, and the distribution of brain states during naturalistic action observation.

## Introduction

Learning from another person’s actions begins before the action is complete. An observer must identify the movement being performed, retain its sequence, and use the unfolding context to anticipate what may come next. These processes make observation an important route to acquiring and refining motor skills (Ramsey et al., 2021). The temporal structure of an action matters: when the same component movements are presented in a meaningful versus scrambled order, sensorimotor and visual responses change, indicating that the brain represents relations between successive acts as well as the acts themselves (Thomas et al., 2018). Predictive accounts of action observation likewise emphasize the use of prior experience to constrain upcoming movement (Kilner et al., 2007). This perspective suggests a broad question for natural behavior: how is an observer’s learned action repertoire expressed in the ongoing brain response to another person’s movements?

Dance offers a powerful way to examine that question. Training creates large differences in the movements people can execute, recognize, and anticipate, while the same performance can be shown to expert and novice observers (Karpati et al., 2015). Dancers viewing movements from their own repertoire show stronger responses in premotor, parietal, and temporal areas than when viewing unfamiliar dance styles (Calvo-Merino et al., 2005, 2006). The observation system is also altered by learning itself: physical rehearsal and observational exposure both change responses to dance sequences (Cross et al., 2006, 2009). At a finer temporal scale, dance expertise modulates alpha/beta activity during action viewing (Orgs et al., 2008). Related work in sport and tango links expertise to improved action anticipation and to neural responses preceding the observed action outcome (Aglioti et al., 2008; Amoruso et al., 2014). Across these studies, experience consistently changes action observation; the remaining challenge is to characterize its expression during a long, uninterrupted stream of movement.

A performance is more than a series of isolated movement clips. The viewer encounters changing relations among performers, alternations between coordinated and independent movement, and new visual contexts that may reshape the meaning of a gesture. Naturalistic fMRI is suited to this setting because it preserves the temporal sequence of events. Intersubject correlation (ISC) reveals when a common stimulus evokes similar responses across viewers (Hasson et al., 2004; Nastase et al., 2019), while studies of continuous narratives show that distributed brain responses change as events unfold and context accumulates (Baldassano et al., 2017; Simony et al., 2016). Coactivation-pattern analysis offers another view of this process by identifying recurring configurations of whole-brain activity (Liu et al., 2013). In dance, the performance itself also supplies measurable temporal features: synchrony and acceleration among performers rise and fall as their interactions develop. Group Study is especially useful because the performance, its choreographic score, and the performers’ movements have been documented (Vicary et al., 2017).

The original fMRI study of Group Study established an important link between movement on stage and shared responses in the audience. Dancers reported greater felt understanding of the performance and showed more similar neural and behavioral responses to one another than novices; performer synchrony predicted spectator ISC in both groups (Orgs et al., 2024). These findings identify a collective response to observed movement and show that expertise matters for how spectators share the experience. Yet two properties of the viewing response remain to be connected to that account. First, similarity between viewers does not describe the temporal organization of each viewer’s regional BOLD signal. Second, a common stimulus can drive recurrent whole-cortex states even when observers differ in the proportion of time they spend in those states. Examining these properties can locate experience effects within individual brain dynamics as well as between observers.

Brain entropy (BEN) provides a measure of the first property. In fMRI, regional BEN is commonly estimated with sample entropy, which evaluates the recurrence of short BOLD signal patterns under specified embedding and tolerance parameters (Richman and Moorman, 2000; Wang et al., 2014). Higher BEN in this framework denotes less regular continuation of those patterns; it does not by itself identify a particular cognitive operation. The measure captures a feature of local temporal organization that differs from activation amplitude, spectral power, and the agreement between viewers assessed by ISC. Our previous movie-watching study showed that regional BEN can be measured reliably during naturalistic viewing and that movie viewing, relative to rest, changes BEN differently across sensory and association cortex (Song and Wang, 2024). In separate work, resting BEN was associated with the magnitude of task activation and deactivation (Lin et al., 2022). These findings make BEN a useful candidate for asking whether learned action experience shapes local BOLD dynamics while people watch the same continuous performance.

We therefore examine dance observation at three connected levels. We first test whether dancers and novices differ in regional BEN across the full film and in successive viewing windows, moving from atlas voxels to parcels and functional networks. We next compare their shared time courses within each experience group and separately assess each spectator’s similarity to the choreographer. Finally, we identify recurring brain states without using group labels and ask whether their occupancy differs by dance experience and whether their occurrence follows measured movement features. This design links local temporal irregularity, collective neural responses, and performance-related state changes within one viewing episode. It tests for neural signatures of prior action experience during observation, while the existing measures of felt understanding and performer movement provide behavioral context for interpreting those signatures.

## Methods

### Participants and stimulus

We reanalyzed the dance-viewing fMRI dataset reported by Orgs et al. (2024). Of the 28 recruited participants, one dancer’s and one novice’s scans were unavailable, leaving 14 professional dancers, 11 novices, and the choreographer. The choreographer was used only as a reference for spectator–choreographer analyses and was excluded from dancer–novice comparisons. Novices had no formal dance training and fewer than five years of casual dance participation.

During scanning, participants watched Group Study, a continuous, single-shot film of ten performers lasting 33 min 54 s. The performance had no musical accompaniment (Vicary et al., 2017; Orgs et al., 2024). Analyses used the first 678 of the 680 acquired BOLD volumes (TR = 3 s), corresponding to the film’s approximately 2034-s duration. Ethical approval and informed consent are detailed in Orgs et al. (2024).

### MRI data acquisition and preprocessing

MRI data were acquired on a Siemens Magnetom Trio 3 T scanner with an eight-channel head coil. The T2*-weighted gradient-echo EPI sequence comprised 41 axial slices (3-mm isotropic voxels; field of view = 192 mm; TR = 3000 ms; TE = 31 ms) and 680 volumes. A 1-mm isotropic T1-weighted image was acquired after the functional run (Orgs et al., 2024).

We obtained the raw BIDS images from OpenNeuro dataset ds004783 (https://openneuro.org/datasets/ds004783/versions/1.0.1) and processed them locally with fMRIPrep 23.1.4, producing BOLD images in MNI152NLin2009cAsym space (Esteban et al., 2019). For BEN, ISC, and CAP analyses, we further processed these images with Nilearn (Nilearn contributors, 2023). The confound matrix included white-matter and cerebrospinal-fluid signals and their derivatives; six motion parameters and their derivatives; standardized DVARS, framewise displacement, and RMS displacement; one temporal CompCor component; and three anatomical CompCor components (Behzadi et al., 2007). Missing confound values were replaced with zero. The BOLD series were detrended, band-pass filtered at 0.009–0.08 Hz, cleaned of the specified confounds, and standardized using the sample standard deviation. We then applied 6-mm FWHM spatial smoothing. Global signal regression was not performed. BEN, ISC, and CAP were calculated from these processed series using the first 678 volumes. ALFF, fALFF, and ReHo were calculated separately from the fMRIPrep BOLD images, as described under Regional BOLD dynamics.

### Regional BOLD dynamics

Static and dynamic BEN. We quantified regional BOLD temporal irregularity using sample entropy (Richman and Moorman, 2000; Wang et al., 2014). Static BEN was calculated at each valid Schaefer atlas voxel from the first 678 BOLD volumes, with embedding dimension (m=3) and tolerance (r=0.6) times that voxel’s time-series standard deviation. We then averaged voxel estimates within each of the 400 parcels and calculated each network value as the equal-weight mean of its constituent parcels.

For dynamic BEN, we first averaged the BOLD signal within each parcel and then calculated sample entropy in overlapping 60-TR (180-s) windows advanced by 10 TR (30 s). This yielded 62 windows per participant. The windowed estimator used (m=3), a tolerance of (0.6) times the within-window standard deviation, and a 0.5 pseudocount for matching-pattern counts. For each parcel, we calculated the mean and standard deviation of BEN across windows; network values were equal-weight means of the corresponding parcel values.

Complementary regional measures. ALFF, fALFF, and ReHo were calculated voxelwise from the first 678 volumes of the fMRIPrep BOLD images after conversion to percent signal change and regression of an intercept, linear trend, and tissue, motion, displacement, DVARS, and CompCor confounds. ALFF was the mean Fourier amplitude in the 0.01–0.08 Hz band. fALFF was the summed amplitude in that band divided by the summed amplitude across all nonzero frequencies up to the Nyquist frequency (Zou et al., 2008). For ReHo, the nuisance-adjusted series were band-pass filtered at 0.01–0.08 Hz, and Kendall’s coefficient of concordance was calculated for each voxel and its valid 3 × 3 × 3 neighborhood (Zang et al., 2004). ReHo estimates required at least 20 valid neighborhood voxels. Each metric was averaged within parcels and then across parcels within networks.

### Shared neural responses and coactivation states

We calculated ISC from the film-aligned parcel BOLD time series. For each parcel, choreographer-paired ISC was the Pearson correlation between a spectator’s time series and the choreographer’s time series, transformed with Fisher’s z. Within-group ISC was calculated for every dancer–dancer pair (91 dyads) and every novice–novice pair (55 dyads). Group differences were tested with 4,999 permutations of participant labels that preserved the 14-dancer/11-novice group sizes (Nastase et al., 2019).

To identify recurring coactivation patterns (CAPs), parcel signals were standardized across time within each spectator. Each 400-parcel frame was then spatially centered and scaled to unit length. Four CAP centroids were fitted to frames pooled from the 25 spectators without group labels, and the choreographer’s frames were subsequently assigned to these centroids (Liu et al., 2013). Fractional occupancy was the proportion of a participant’s 678 frames assigned to each state.

Performer synchrony and acceleration were derived from wrist accelerometry, and visual change was measured from successive film frames (Orgs et al., 2024). We aligned these features to the BOLD series using the identified film-series offset and a 5-s hemodynamic delay. Complete feature data were available for 57 of the 62 dynamic analysis windows.

### Statistical analysis

For each parcel and network, dancer–novice differences in static BEN, dynamic BEN, ALFF, fALFF, and ReHo were estimated with linear models containing group, regional GMV, age, sex, and mean framewise displacement. Choreographer-paired ISC models contained group, age, sex, and mean framewise displacement. Within-group ISC differences were evaluated with participant-label permutation tests. CAP occupancy was compared between groups using models adjusted for age, sex, and mean framewise displacement. Associations between film features and dynamic brain measures were estimated within participants and then summarized across spectators.

*P* values were corrected for multiple comparisons using the Benjamini–Hochberg false discovery rate (FDR) procedure (Benjamini and Hochberg, 1995). Correction was applied across the 400 parcels and, separately, across the seven networks for each measure; the four CAP occupancy tests were corrected together. All tests were two-sided, with FDR-adjusted q<0.05 defining statistical significance.

## Results

### Dance experience is expressed in local BOLD temporal irregularity

Dancers had higher static BOLD sample entropy than novices in 32 of 400 parcels after adjustment for regional GMV, age, sex, and mean framewise displacement. The effect extended across visual (q = .0438), somatomotor (q = .0201), dorsal-attention (q = .0201), salience/ventral-attention (q = .0201), control (q = .0201), and default-mode (q = .0466) networks. The limbic network did not meet the seven-network threshold (q = .475; Figure 1). Adding total GMV to the local-GMV model retained corrected effects in somatomotor, dorsal-attention, salience/ventral-attention, and control networks, with 38 significant parcels. Thus, the principal local result spans sensory, action-related, and association systems rather than a single action-observation region.

**Figure 1.**
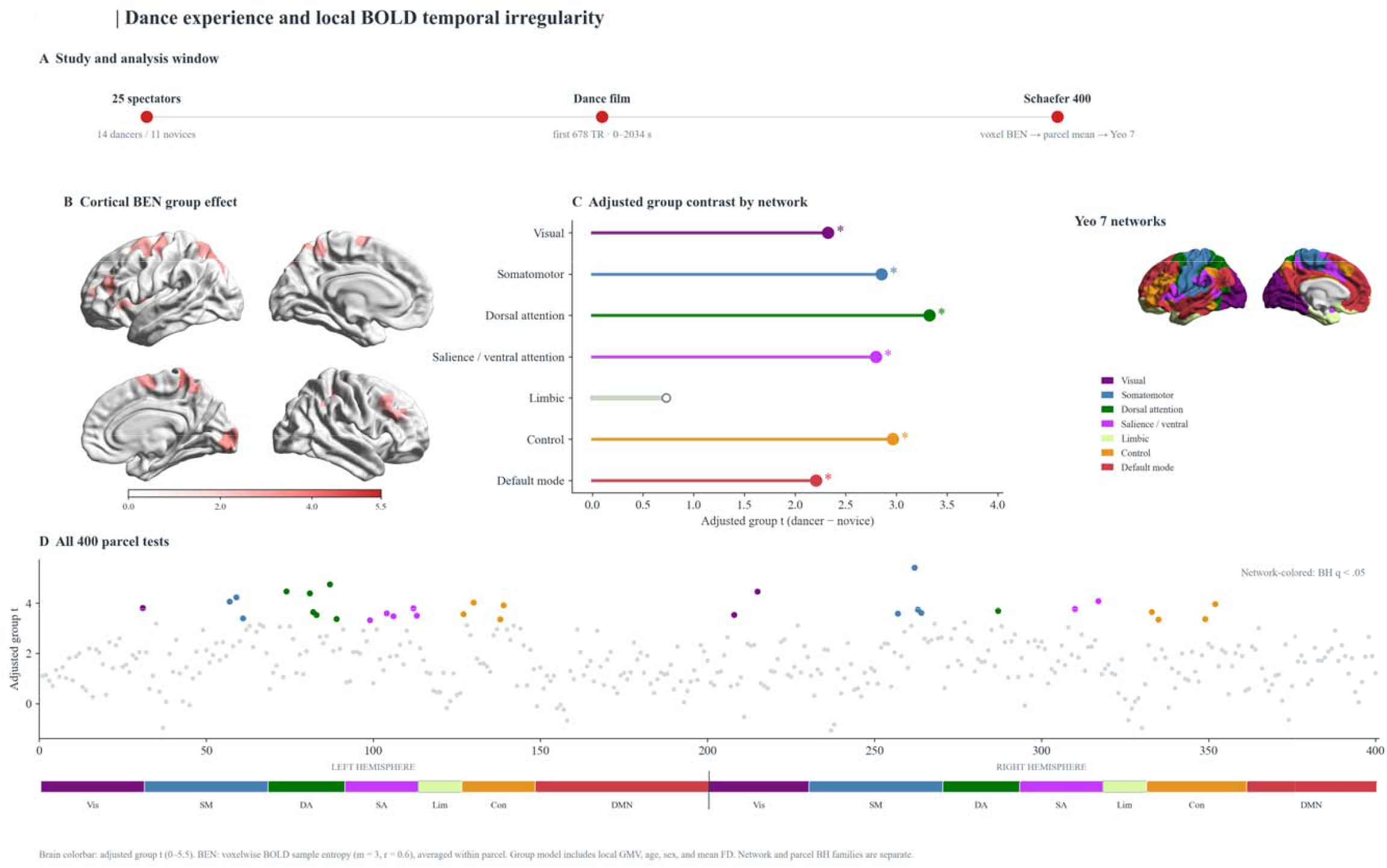
Static BEN. Voxelwise BOLD sample entropy was averaged into 400 parcels and seven networks. Dancer–novice models include local GMV, age, sex, and mean framewise displacement. Filled network-color points mark Benjami Hochberg q < .05; 32 parcels and six networks passed their separate spatial correction families.

### The dynamic effect concentrates in control and default networks

Across 62 windows, mean parcel-based BOLD entropy was higher in dancers in the control and default-mode networks (q = .0384 for each; Figure 2). This result places the group effect within the continuing performance, using a time-resolved measure with a different signal-aggregation step from static voxel BEN. At the 400-parcel level, dynamic entropy did not pass correction, and neither across-window entropy variability nor group differences in movement-feature slopes yielded corrected network effects.

**Figure 2.**
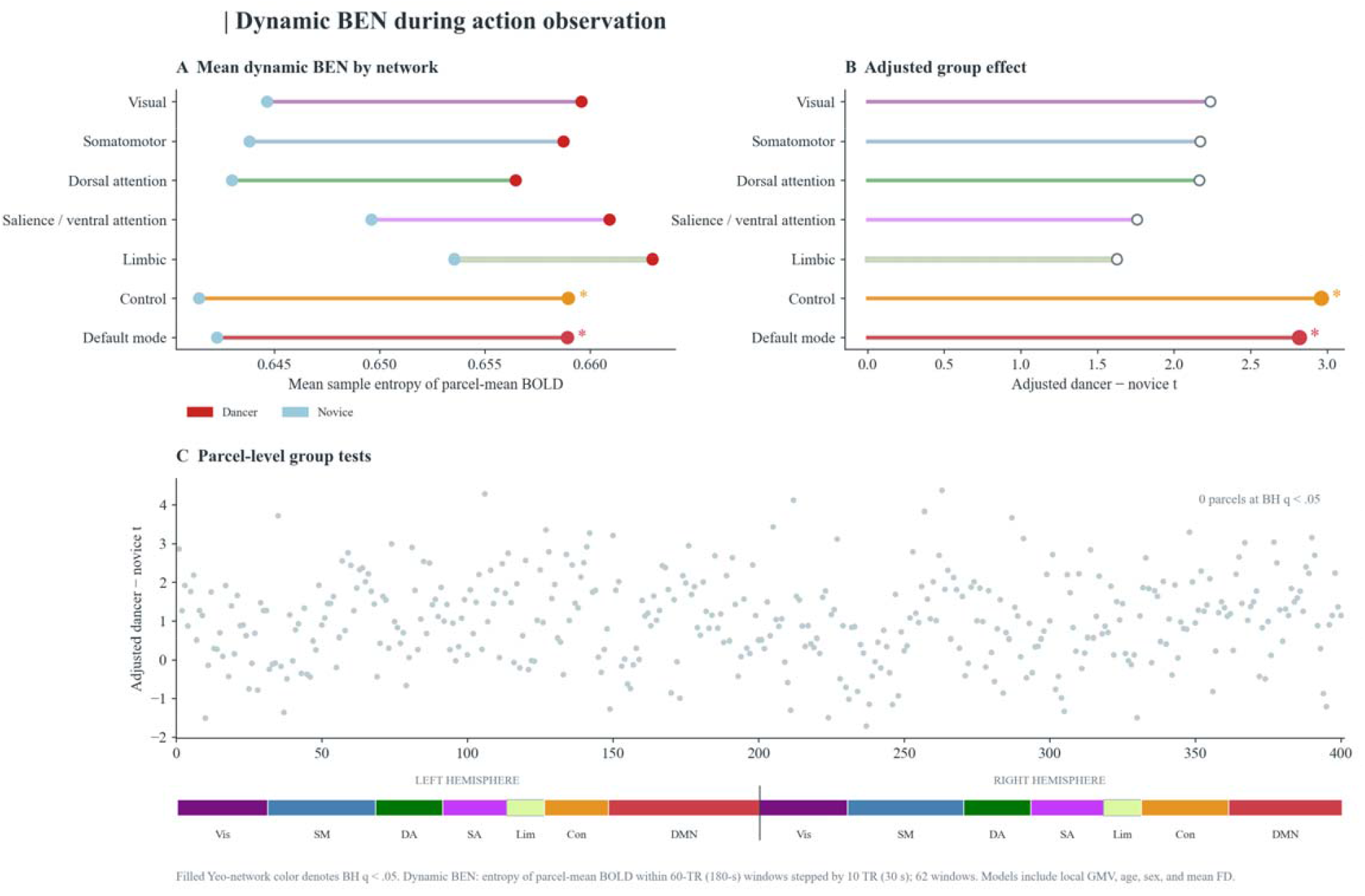
Dynamic BEN. Entropy of parcel-mean BOLD was calculated in 60-TR windows stepped by 10 TR. Mean dynamic BEN was higher in dancers in control and default-mode networks after adjustment; no parcel p colors follow the Yeo assignment.

### Shared spectator responses carry a complementary expertise signature

Dancer-dancer ISC exceeded novice-novice ISC in somatomotor (difference in Fisher z = .0267), dorsal-attention (.0167), salience/ventral-attention (.0224), control (.0149), and default-mode (.0193) networks (q = .0423 for each; Figure 3). Visual-network ISC was in the same direction (q = .0590); no individual parcel passed the 400-parcel correction. The network-level result supports the original report of stronger within-group synchrony among dancers (Orgs et al., 2024) under the present 678-volume, atlas-based analysis. In a covariate-adjusted sensitivity analysis, the salience/ventral-attention effect remained corrected. Choreographer-paired ISC did not show a corrected group difference at parcel or network level, and static BEN was not associated with choreographer-paired ISC after correction. The shared-response evidence is therefore strongest for similarity among trained spectators, not similarity to the reference individual.

**Figure 3.**
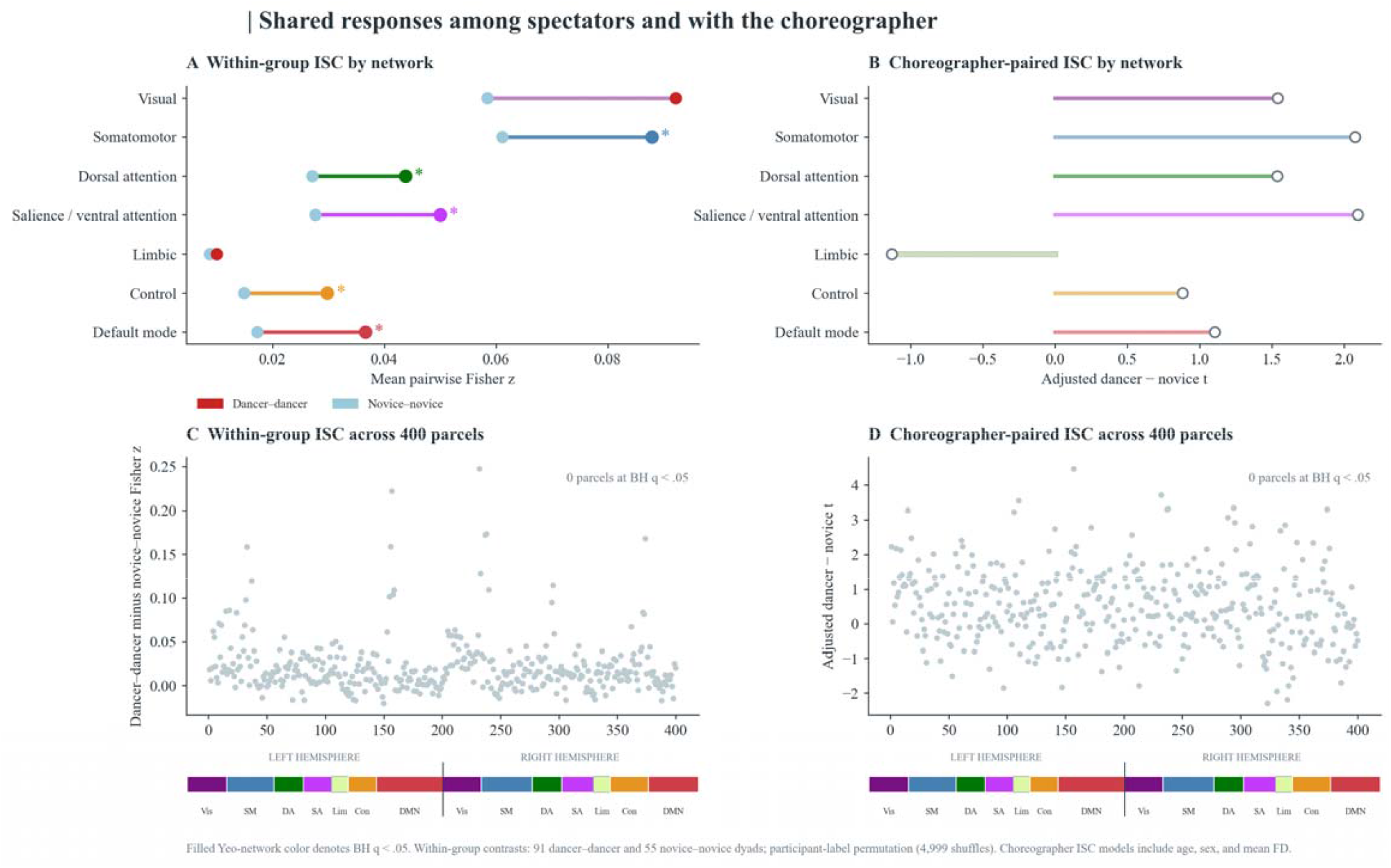
Shared responses. Within-group ISC compares 91 dancer–dancer with 55 novice–novice dyads using participant-label permutation. Five networks passed seven-network BH correction. Choreographer-paired ISC sh difference at network or parcel level.

### Film movement is reflected in recurring audience brain states

Four group-blind CAPs partitioned the 16,950 spectator frames. Dancers occupied State 1 less often than novices (23.35% versus 26.71%; adjusted group coefficient = -.0289, q = .0103) and State 3 more often (27.74% versus 24.05%; coefficient = .0347, q = .0103; Figure 4). Performer synchrony and acceleration were positively associated with State 1 occurrence across spectators (q = .0063 and .0151) and negatively associated with State 2 occurrence (q = .0063 and .0059). The film-feature slopes did not show corrected dancer-novice interactions. Together, the occupancy and feature results locate experience-related differences in the distribution of common states while identifying movement features that accompany state transitions across the full audience.

**Figure 4.**
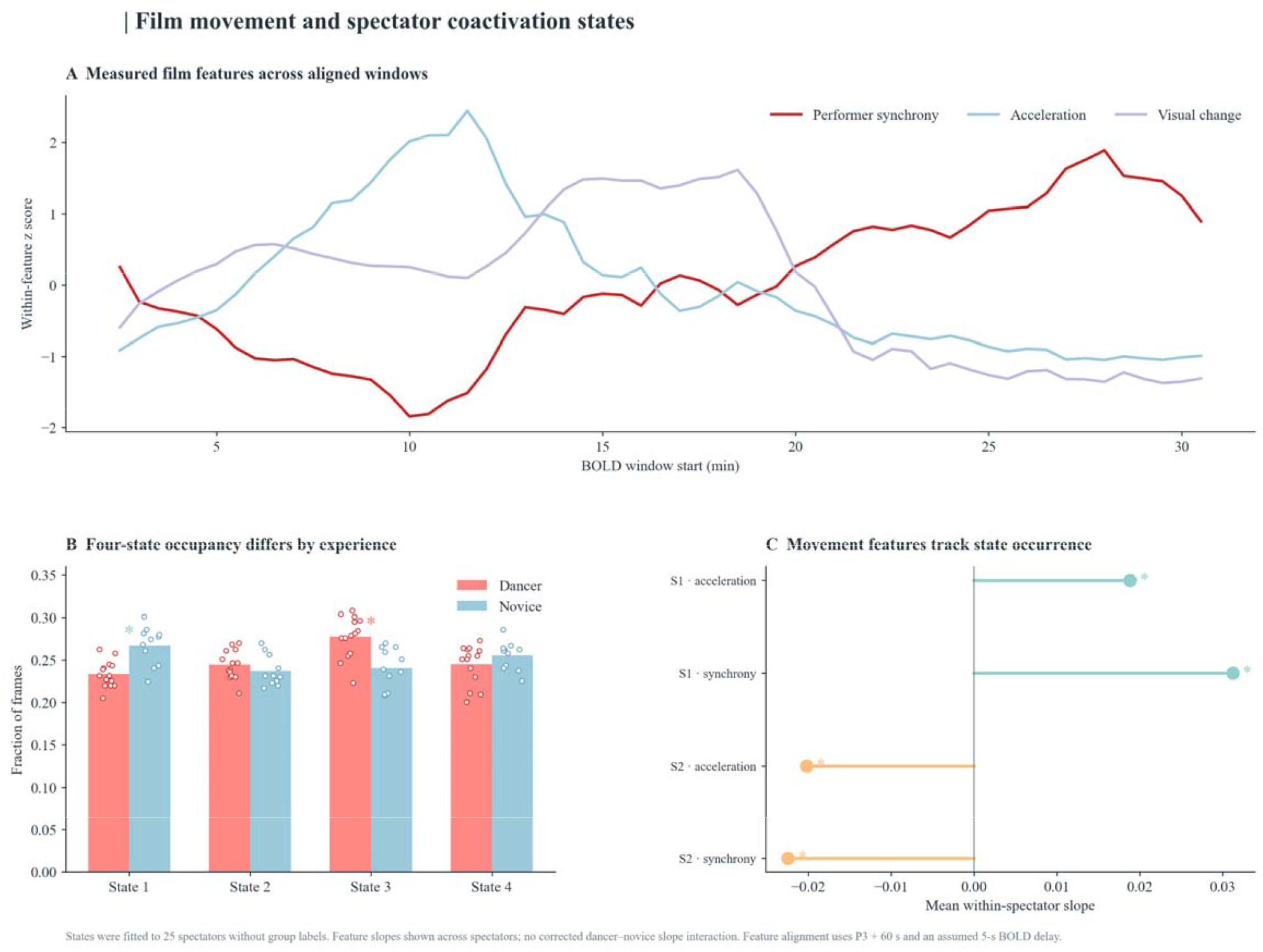
Film movement and brain states. Four CAPs were fitted without group labels. Dancers occupied State 1 less and State 3 more than novices. Performer synchrony and acceleration tracked State 1 positively and State 2 negatively ac group-by-feature interactions were absent.

### Complementary local measures

The spectral comparison yielded lower fALFF in dancers in eight parcels and six networks: somatomotor, dorsal attention, salience/ventral attention, limbic, control, and default mode (network q = .0465; Supplementary Figure 1). ALFF and ReHo had no corrected parcel or network group effects. fALFF therefore adds a spectral-composition result to the local temporal-irregularity result without reproducing the same contrast across every local measure. The corrected network findings are summarized in Table 1.

**Table 1.** Corrected network-level findings.

| **Endpoint** | **Networks** | **Dancer vs novice** | **BH q** |
|---|---|---|---|
| Static BEN | Visual | Higher | .0438 |
| Static BEN | Somatomotor, dorsal attention, salience/ventral attention, control | Higher | .0201 |
| Static BEN | Default mode | Higher | .0466 |
| Dynamic BEN mean | Control, default mode | Higher | .0384 |
| Within-group ISC | Somatomotor, dorsal attention, salience/ventral attention, control, default mode | Dancer dyads higher | .0423 |
| fALFF | Six nonvisual networks | Lower | .0465 |

Note. Static and dynamic BEN, within-group ISC, and fALFF are distinct endpoints. q values are corrected within the named seven-network family. fALFF is a supplementary local measure.

## Discussion

Learning from another person’s actions depends on following movement over time. An observer sees what a performer is doing now while using previous movements and prior experience to interpret how the action is developing. Dance makes this process especially visible: expertise changes responses to familiar movements and to sequences learned through practice or observation (Calvo-Merino et al., 2005, 2006; Cross et al., 2006, 2009). During an uninterrupted performance, those learned representations can be engaged repeatedly as performers change speed, coordinate with one another, and move into new configurations. Our results show that dance experience is expressed in regional BOLD temporal irregularity, similarity between spectators, and the occupancy of recurring brain states during that unfolding observation.

Our previous movie-viewing study provides a foundation for interpreting the BEN findings. Song and Wang (2024) showed that regional BEN can be measured reliably during movie viewing and that sustained viewing changes BEN across sensory and association cortex. The present results bring action experience into this naturalistic framework. Dancers and novices watched the same film, yet dancers showed higher static BEN across 32 parcels and six networks. Mean windowed BEN was also higher in dancers in control and default-mode networks. Sample entropy describes the recurrence of short BOLD patterns: these findings indicate that prior dance experience is associated with how regularly regional activity evolves during observation, both across the full performance and in successive viewing periods.

The network distribution fits a view of action observation that reaches beyond the recognition of individual movements. In tango, expertise influences anticipatory neural activity and the interpretation of an action in context (Amoruso et al., 2014). Work on observational learning likewise places sensorimotor processing alongside executive and social processes that support learning from others’ actions (Ramsey et al., 2021). A long performance continually supplies new context for interpreting movement. Naturalistic studies have shown that higher-order cortical responses depend on information accumulated over time and change as meaningful sequences unfold (Hasson et al., 2008; Baldassano et al., 2017; Simony et al., 2016). The control- and default-mode windowed BEN effects are therefore particularly relevant to the sustained demands of following the choreography, while the broader static effect shows that experience is expressed across multiple systems engaged by the performance.

Experience also shaped what spectators shared. Dancer–dancer ISC exceeded novice–novice ISC in five networks, extending the original report of stronger neural agreement among trained spectators (Orgs et al., 2024). The dancers were scanned separately, so this agreement reflects more similar timing of responses to the same evolving film. BEN and ISC describe complementary aspects of that response: BEN characterizes temporal patterning within a spectator’s regional signal, whereas ISC measures correspondence between spectators’ signals. The choreographer-paired comparison addresses similarity to one individual’s response; the strongest expertise effect in this dataset emerged instead from the shared responses of trained viewers.

The CAP results connect those audience responses to events on stage. Dancers spent less time in State 1 and more time in State 3 than novices, although the four states were identified without group labels. Performer synchrony and acceleration were associated with State 1 and State 2 occurrence across the audience. Brain-state dynamics during other films have likewise been linked to features of the unfolding stimulus (van der Meer et al., 2020). In Group Study, measurable properties of performers’ movement were reflected in the occurrence of common audience states, and dance experience was reflected in how spectators distributed their time among those states. This places the expertise effect within a brain response that remains engaged with the changing performance.

The complementary measures specify the regional signal differences further. Dancers showed lower fALFF in six networks, indicating a smaller proportion of spectral amplitude in the 0.01–0.08 Hz band. ALFF and ReHo showed no corrected group differences. Together, BEN and fALFF show that dance experience was associated with temporal pattern recurrence and spectral composition during viewing. The BEN result adds to the movie-viewing evidence of Song and Wang (2024) by showing how an observer’s acquired action repertoire is reflected in regional BOLD dynamics under a shared naturalistic stimulus.

## Conclusion

Dance experience was expressed at several levels of the response to an uninterrupted performance: higher regional BOLD sample entropy, greater neural agreement among trained spectators, and altered occupancy of recurring brain states. Performer synchrony and acceleration were associated with the occurrence of those states across the audience. These findings connect prior action experience with the temporal organization of individual and shared brain responses during naturalistic observation.

---

**Supplementary Figure 1.**
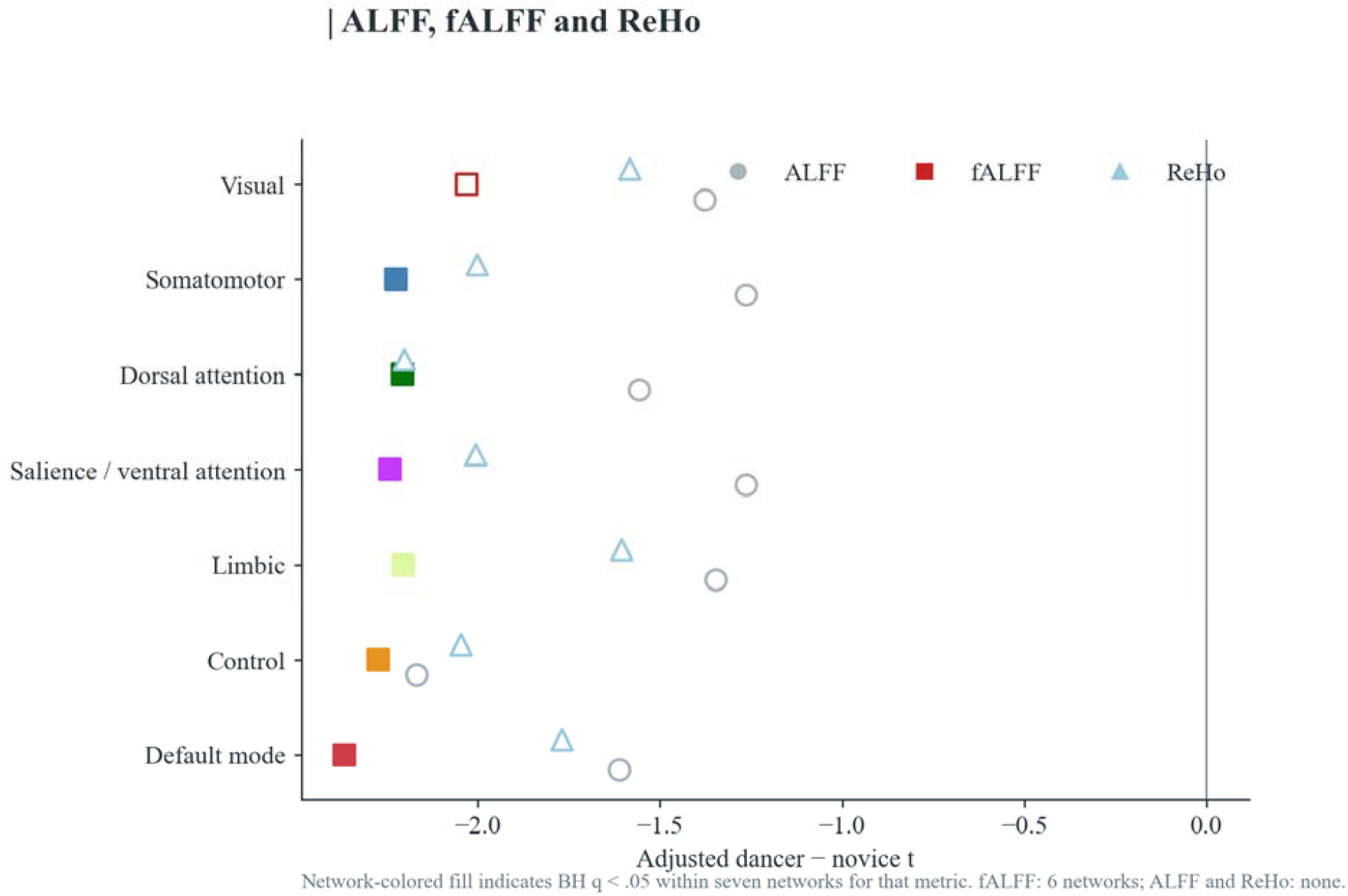
Other local BOLD measures. fALFF was lower in dancers in six networks; ALFF and ReHo had no corrected network group differences. Each metric used its own seven-network BH family.

## References

Aglioti, S. M., Cesari, P., Romani, M., & Urgesi, C. (2008). Action anticipation and motor resonance in elite basketball players. Nature Neuroscience, 11, 1109–1116. 10.1038/nn.2182

Amoruso, L., Sedeño, L., Huepe, D., et al. (2014). Time to tango: Expertise and contextual anticipation during action observation. NeuroImage, 98, 366–385. 10.1016/j.neuroimage.2014.05.005

Baldassano, C., Chen, J., Zadbood, A., Pillow, J. W., Hasson, U., & Norman, K. A. (2017). Discovering event structure in continuous narrative perception and memory. Neuron, 95, 709-721.e5. 10.1016/j.neuron.2017.06.041

Behzadi, Y., Restom, K., Liau, J., & Liu, T. T. (2007). A component based noise correction method (CompCor) for BOLD and perfusion based fMRI. NeuroImage, 37, 90–101. 10.1016/j.neuroimage.2007.04.042

Benjamini, Y., & Hochberg, Y. (1995). Controlling the false discovery rate: A practical and powerful approach to multiple testing. Journal of the Royal Statistical Society: Series B (Methodological), 57, 289–300. 10.1111/j.2517-6161.1995.tb02031.x

Calvo-Merino, B., Glaser, D. E., Grèzes, J., Passingham, R. E., & Haggard, P. (2005). Action observation and acquired motor skills: An fMRI study with expert dancers. Cerebral Cortex, 15, 1243–1249. 10.1093/cercor/bhi007

Calvo-Merino, B., Grèzes, J., Glaser, D. E., Passingham, R. E., & Haggard, P. (2006). Seeing or doing? Influence of visual and motor familiarity in action observation. Current Biology, 16, 1905–1910. 10.1016/j.cub.2006.07.065

Cross, E. S., Hamilton, A. F. de C., & Grafton, S. T. (2006). Building a motor simulation de novo: Observation of dance by dancers. NeuroImage, 31, 1257–1267. 10.1016/j.neuroimage.2006.01.033

Cross, E. S., Kraemer, D. J. M., Hamilton, A. F. de C., Kelley, W. M., & Grafton, S. T. (2009). Sensitivity of the action observation network to physical and observational learning. Cerebral Cortex, 19, 315–326. 10.1093/cercor/bhn083

Esteban, O., Markiewicz, C. J., Blair, R. W., et al. (2019). fMRIPrep: A robust preprocessing pipeline for functional MRI. Nature Methods, 16, 111–116. 10.1038/s41592-018-0235-4

Hasson, U., Nir, Y., Levy, I., Fuhrmann, G., & Malach, R. (2004). Intersubject synchronization of cortical activity during natural vision. Science, 303, 1634–1640. 10.1126/science.1089506

Hasson, U., Yang, E., Vallines, I., Heeger, D. J., & Rubin, N. (2008). A hierarchy of temporal receptive windows in human cortex. Journal of Neuroscience, 28, 2539–2550. 10.1523/JNEUROSCI.5487-07.2008

Karpati, F. J., Giacosa, C., Foster, N. E. V., Penhune, V. B., & Hyde, K. L. (2015). Dance and the brain: A review. Annals of the New York Academy of Sciences, 1337, 140–146. 10.1111/nyas.12632

Kilner, J. M., Friston, K. J., & Frith, C. D. (2007). Predictive coding: An account of the mirror neuron system. Cognitive Processing, 8, 159–166. 10.1007/s10339-007-0170-2

Lin, L., Chang, D., Song, D., Li, Y., & Wang, Z. (2022). Lower resting brain entropy is associated with stronger task activation and deactivation. NeuroImage, 249, 118875. 10.1016/j.neuroimage.2022.118875

Liu, X., Chang, C., & Duyn, J. H. (2013). Decomposition of spontaneous brain activity into distinct fMRI co-activation patterns. Frontiers in Systems Neuroscience, 7, 101. 10.3389/fnsys.2013.00101

Nastase, S. A., Gazzola, V., Hasson, U., & Keysers, C. (2019). Measuring shared responses across subjects using intersubject correlation. Social Cognitive and Affective Neuroscience, 14, 667–685. 10.1093/scan/nsz037

Nilearn contributors. (2023). Nilearn [Computer software]. 10.5281/zenodo.8397156

Orgs, G., Dombrowski, J.-H., Heil, M., & Jansen-Osmann, P. (2008). Expertise in dance modulates alpha/beta event-related desynchronization during action observation. European Journal of Neuroscience, 27, 3380–3384. 10.1111/j.1460-9568.2008.06271.x

Orgs, G., Vicary, S., Sperling, M., Richardson, D. C., & Williams, A. L. (2024). Movement synchrony among dance performers predicts brain synchrony among dance spectators. Scientific Reports, 14, 22079. 10.1038/s41598-024-73438-0

Ramsey, R., Kaplan, D. M., & Cross, E. S. (2021). Watch and learn: The cognitive neuroscience of learning from others’ actions. Trends in Neurosciences, 44, 478–491. 10.1016/j.tins.2021.01.007

Richman, J. S., & Moorman, J. R. (2000). Physiological time-series analysis using approximate entropy and sample entropy. American Journal of Physiology Heart and Circulatory Physiology, 278, H2039–H2049. 10.1152/ajpheart.2000.278.6.H2039

Schaefer, A., Kong, R., Gordon, E. M., et al. (2018). Local-global parcellation of the human cerebral cortex from intrinsic functional connectivity MRI. Cerebral Cortex, 28, 3095–3114. 10.1093/cercor/bhx179

Simony, E., Honey, C. J., Chen, J., et al. (2016). Dynamic reconfiguration of the default mode network during narrative comprehension. Nature Communications, 7, 12141. 10.1038/ncomms12141

Song, D.-H., & Wang, Z. (2024). Regional brain entropy during movie-watching [Preprint]. bioRxiv. 10.1101/2024.06.12.598767

Thomas, R. M., De Sanctis, T., Gazzola, V., & Keysers, C. (2018). Where and how our brain represents the temporal structure of observed action. NeuroImage, 183, 677–697. 10.1016/j.neuroimage.2018.08.056

van der Meer, J. N., Breakspear, M., Chang, L. J., Sonkusare, S., & Cocchi, L. (2020). Movie viewing elicits rich and reliable brain state dynamics. Nature Communications, 11, 5004. 10.1038/s41467-020-18717-w

Vicary, S., Sperling, M., von Zimmermann, J., Richardson, D. C., & Orgs, G. (2017). Joint action aesthetics. PLOS ONE, 12, e0180101. 10.1371/journal.pone.0180101

Wang, Y., Zhao, Q., Ji, Q., Jin, X., Zhou, C., & Lu, Y. (2024). fMRI evidence of movement familiarization effects on recognition memory in professional dancers. Cerebral Cortex, 34, bhad490. 10.1093/cercor/bhad490

Wang, Z., Li, Y., Childress, A. R., & Detre, J. A. (2014). Brain entropy mapping using fMRI. PLOS ONE, 9, e89948. 10.1371/journal.pone.0089948

Yeo, B. T. T., Krienen, F. M., Sepulcre, J., et al. (2011). The organization of the human cerebral cortex estimated by intrinsic functional connectivity. Journal of Neurophysiology, 106, 1125–1165. 10.1152/jn.00338.2011

Zang, Y., Jiang, T., Lu, Y., He, Y., & Tian, L. (2004). Regional homogeneity approach to fMRI data analysis. NeuroImage, 22, 394–400. 10.1016/j.neuroimage.2003.12.030

Zou, Q.-H., Zhu, C.-Z., Yang, Y., et al. (2008). An improved approach to detection of amplitude of low-frequency fluctuation (ALFF) for resting-state fMRI: Fractional ALFF. Journal of Neuroscience Methods, 172, 137–141. 10.1016/j.jneumeth.2008.04.012

